# Unique molecular identifier-based single strand mitochondrial DNA sequencing for rare substitution mutations

**DOI:** 10.1101/2025.06.05.657263

**Authors:** Zeshuo E.S. Li, Rachel Dunn, Loïc C. Caloren, Adam S. Ziada, Hailey Chapman, Izabelle Gadawska, Hélène C.F. Côté

## Abstract

The study of mitochondrial genetics has long been limited to polymorphisms and high frequency mutations owing in part to technical and technological limitations in reliably detecting and quantifying rare somatic mutations. Over the past decade or so, the study of rare somatic mitochondrial DNA (mtDNA) variants has expanded and continues to garner increasing interest in a wide range of research fields. Here, we describe a unique molecular identifier (UMI)-based library preparation and Next Generation Sequencing (NGS) method for the identification and quantification of ultra-rare mutations in the mtDNA control region. Our method exploits degenerate primers to label individual mtDNA molecules. This is followed by several purification, quantification and amplification steps, to obtain high quality amplicons for sequencing on the Illumina MiSeq platform. This method enables the use of total genomic DNA extract as starting point for the assay, overcoming the need for organelle isolation and/or mtDNA enrichment, hence broadening the type of specimen that can be studied, while offering cost and time benefits. The assay described herein has been demonstrated to reliably measure somatic variants present at approximately 0.05% variant allele frequency in a variety of tissues, including fresh and frozen biobanked specimens. Using this protocol, library preparation of 300 specimens can be completed by a single individual with general nucleic acid handling experience in approximately 20 days. Given its flexibility and scalability, this method is particularly well suited for epidemiological studies using a large number of specimens.

## Introduction

The mitochondrial genome is circular and comprises 16,569 bp that encodes 13 polypeptides essential for oxidative phosphorylation, 22 tRNA, and 2 rRNA. It also contains a non-coding mitochondrial control region (mCR). Despite advances in sequencing technology, in comparison to nuclear DNA genomics, the study of mitochondrial genomics remains challenging due to the mitochondrial genome’s heterogeneity, high copy number, and high turnover rate^1^.

Increased mitochondrial DNA (mtDNA) mutations, oxidative damage, and dysregulation of mitochondrial fusion and fission are common characteristics of many age related pathologies ^2–4^. Compared to its nuclear homolog, mtDNA is more susceptible to *de novo* mutations due to the limited proofreading ability of the polymerase gamma (Pol ɣ) responsible for its replication and maintenance, and the high oxidative environment within the mitochondria^5,6^. Although considerable progress has been made in characterizing mtDNA mutations and their possible role in mitochondrial dysfunction and disease, studies are often limited to high frequency mtDNA variants, and few have aimed to identify or quantify rare mtDNA mutations. This is in part due to the paucity of robust sequencing methods tailored specifically to mtDNA. Further, some questions are increasingly being raised around the prevalence and reliability of low frequency (1-2% variant allele) mtDNA heteroplasmy^7,8^. Taken together this highlights the need for reliable methods for the detection and quantification of low-level mutations in mtDNA.

It is well recognized that the non-coding mCR sequence is more variable than other regions of the mtDNA that likely have greater functional constraints. Within the mCR, three short hypervariable segments (HVS) are especially likely to harbor mtDNA polymorphisms^9–11^. The high variability at these locations, DNA contamination, and index hopping, are issues that often plague next generation sequencing (NGS), and may artificially inflate the rate of mtDNA mutations reported^12^. Indeed, the debates over what may be an artifactual versus a true mtDNA mutation has been a concern in the field since the days of Sanger sequencing. While NGS has accelerated research, these challenges remain barriers to our understanding of mitochondrial genetics^13^. In recent studies using massively parallel sequencing, the lower limit of mtDNA mutation detection is a variant allele frequency of approximately 1%, as accuracy below this frequency is tenuous^13–15^. However, unique molecular identifier (UMI, see Table 1 for term definitions) or barcoding approaches to mtDNA sequencing allow the quantification and characterization of the largely overlooked less frequent mutations that may play an important role in mitochondrial function.

**Table 1:**
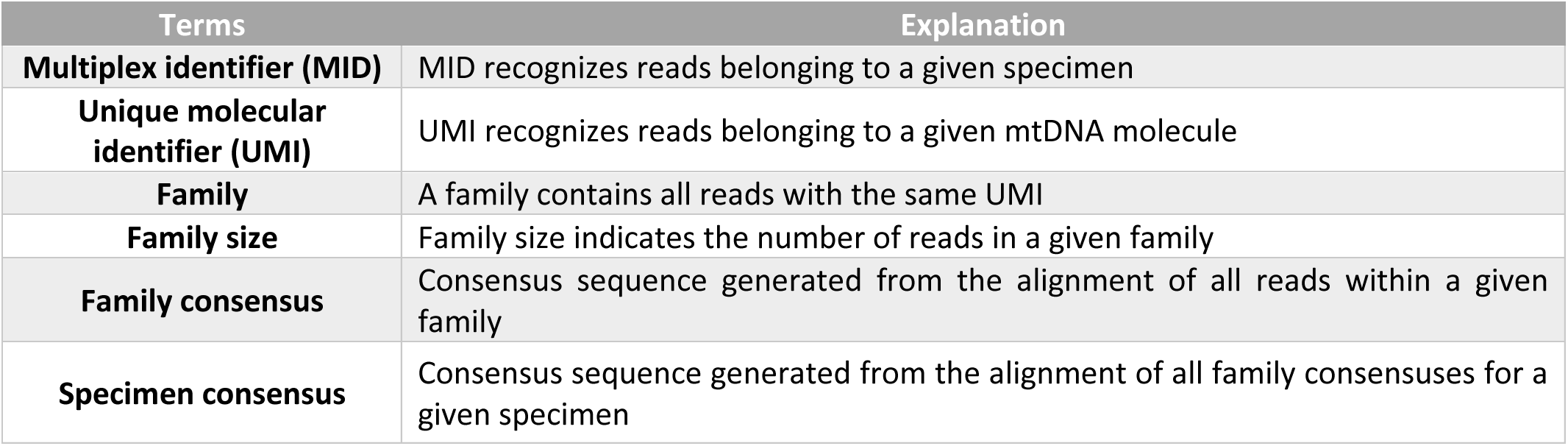
Lexicon of terminology.

### MtDNA sequencing by UMI

The use of UMI, also referred to as primer ID, to identify rare mutations is not new, and has been increasingly used over the past decade. The approach exploits degenerate primer tags to uniquely label individual molecules of mtDNA. This later allows the reconstruction of individual consensus sequences from the original DNA, while filtering out PCR and/or sequencing errors. UMI-based sequencing has been used to detect low-level variants in various viruses^16,17^, identify somatic mutations in tumours^18^, and mosaicism of mutations contributing to disease^19^. Sequencing platforms have developed modified workflows to detect these low-level mutations such as Illumina SafeSeq or Roche CIRCLE-Seq, and other groups have developed methods like duplex sequencing ^14,20^, or SiMSen-Seq^21^. Here, we describe a UMI-based procedure developed for the sequencing of a short segment within the mCR of mtDNA. Figure 1 presents a schematic overview of the methodology.

**Figure 1:**
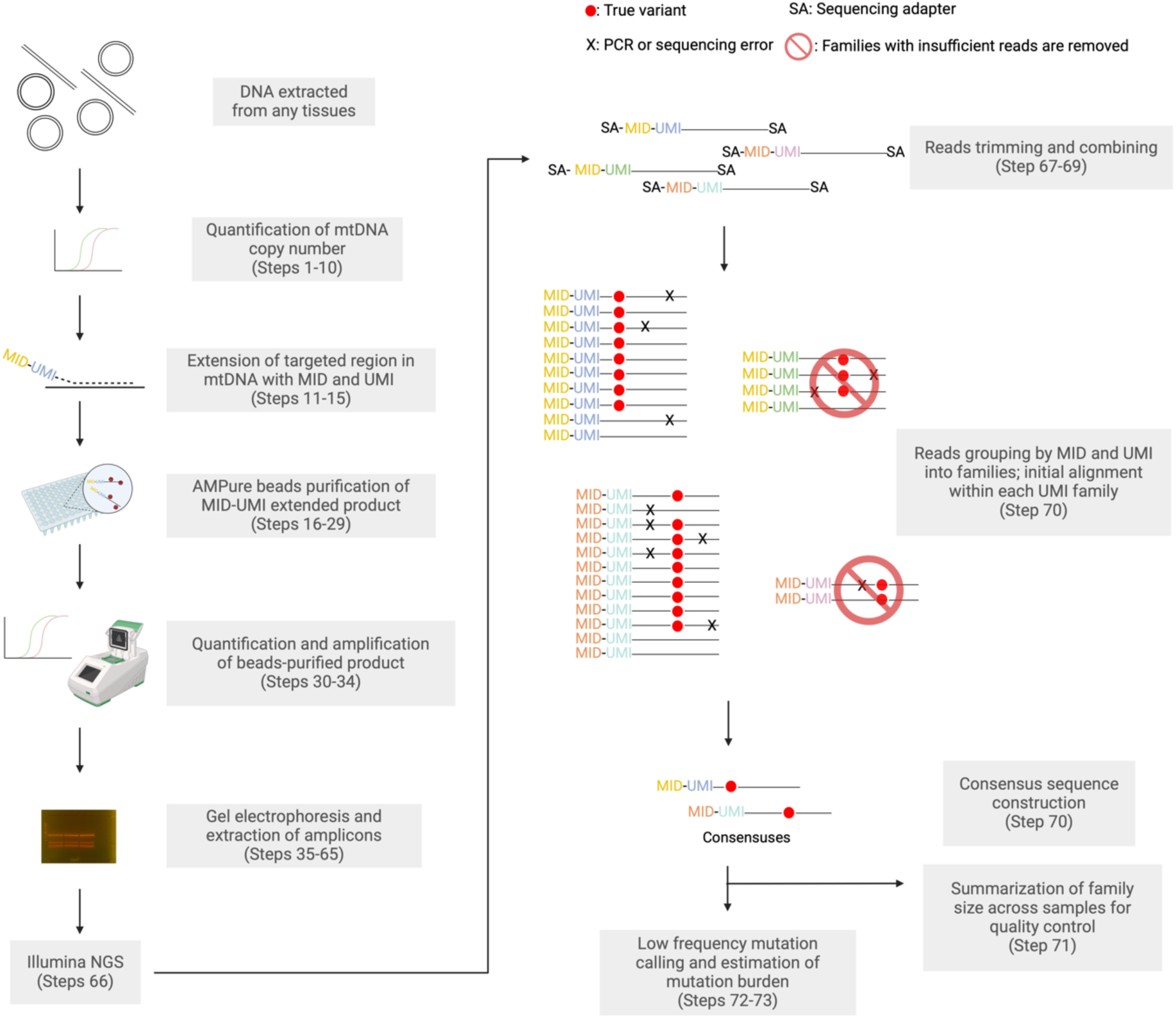
A schematic overview of UMI-based mtDNA sequencing procedure. MtDNA copy number is first quantified in the total DNA extracted from any specimen, a preliminary step for library construction. A primer containing a multiplex identifier (MID) and a unique molecular identifier (UMI) is then used to extend a known number of mtDNA molecules. The resulting tagged product then undergoes several procedures (purification, PCR amplification, and gel electrophoresis), to prepare an amplicon for next generation sequencing (NGS). Subsequent bioinformatics analysis of the NGS data includes trimming sequencing adapters from the reads, merging paired reads, and grouping the reads into families, based on their MID and UMI. Within each UMI family, a consensus sequence is generated through initial alignment. Within each specimen, a specimen consensus sequence is generated through alignment of all family consensus sequences for a given specimen. Mutations are then identified based on the alignment of the family consensuses against the specimen consensus, allowing for the estimation of mutation burden. The distinct sequences of MID and UMI are demarcated by different colors.

The majority of existing UMI assays are aimed at nuclear DNA and do not consider the aforementioned challenges specific to mtDNA. An important consideration when studying mtDNA is that, given the size difference between the two genomes, the amount of nuclear DNA vastly exceeds the amount of mtDNA. Overcoming this issue would historically necessitate isolation of whole mitochondria from fresh tissue, or enrichment for mtDNA. Among the UMI-based protocols for mtDNA sequencing published to date, either organelle isolation, or targeted DNA capture are used to isolate mtDNA^20,22^, steps that require either fresh tissue with minimal cellular damage, or large amounts of DNA, respectively. For the majority of large human research studies using archived frozen specimens, neither of those are feasible. Although large biobanks are increasingly used as a research tool, a lag exists in the development of methodologies that can make the most of these valuable resources, including tailored techniques. Biobanks often collect specimens over years if not decades, giving rise to variability in the quality, quantity, age, and handling of these tissues. All these factors can affect both the yield and the quality of DNA available, which in turn influences the feasibility of processing methods such as mtDNA enrichment. In our development of UMI-based sequencing method, close consideration was paid to factors such as DNA quality and quantity requirements, and labour intensity, to maximize cost-effectiveness and applicability. Our methodology uses total DNA extract as a starting point for mtDNA sequencing library preparation which circumvents the limitations cited above. Our mtDNA sequencing protocol has proven robust and reliable for DNA from biobanked whole blood^23^, and our group has since applied it to several other human tissues, including skeletal muscle biopsy, placenta tissue, cultured fibroblasts, human embryonic stem cell lines, and buffy coat (unpublished data).

### Advantages

Our UMI method successfully removes inter-run specimen contamination and sequencing artefacts. With conventional NGS of mtDNA, we previously observed high mutation signals for a plasmid containing the mtDNA mCR region, our clonal negative control (Figure 2A). These artificially introduced variants were detected at several hotspot positions that are known to be highly variable and often heteroplasmic in the general population. This can be visualized in the mutation patterns of the human whole blood specimens used for our assay high and low mutation controls (Figure 2C-F unpublished data). Further, several of these commonly mutated positions are also the site of documented polymorphisms based on GenBank sequences (MITOMAP: A human Mitochondrial Genome Database^24^), suggesting that these artefacts are likely due to index hopping^25,26^. However, when UMI filtering was applied, artefactual signals observed in the clonal negative control specimen were removed (Figure 2B), and the number of mutations observed in the low mutation controls decreased while the high mutation control maintained its higher burden of mutations (Figure 2D,F). This indicates the successful removal of contamination between sequenced specimens using our UMI methodology.

**Figure 2:**
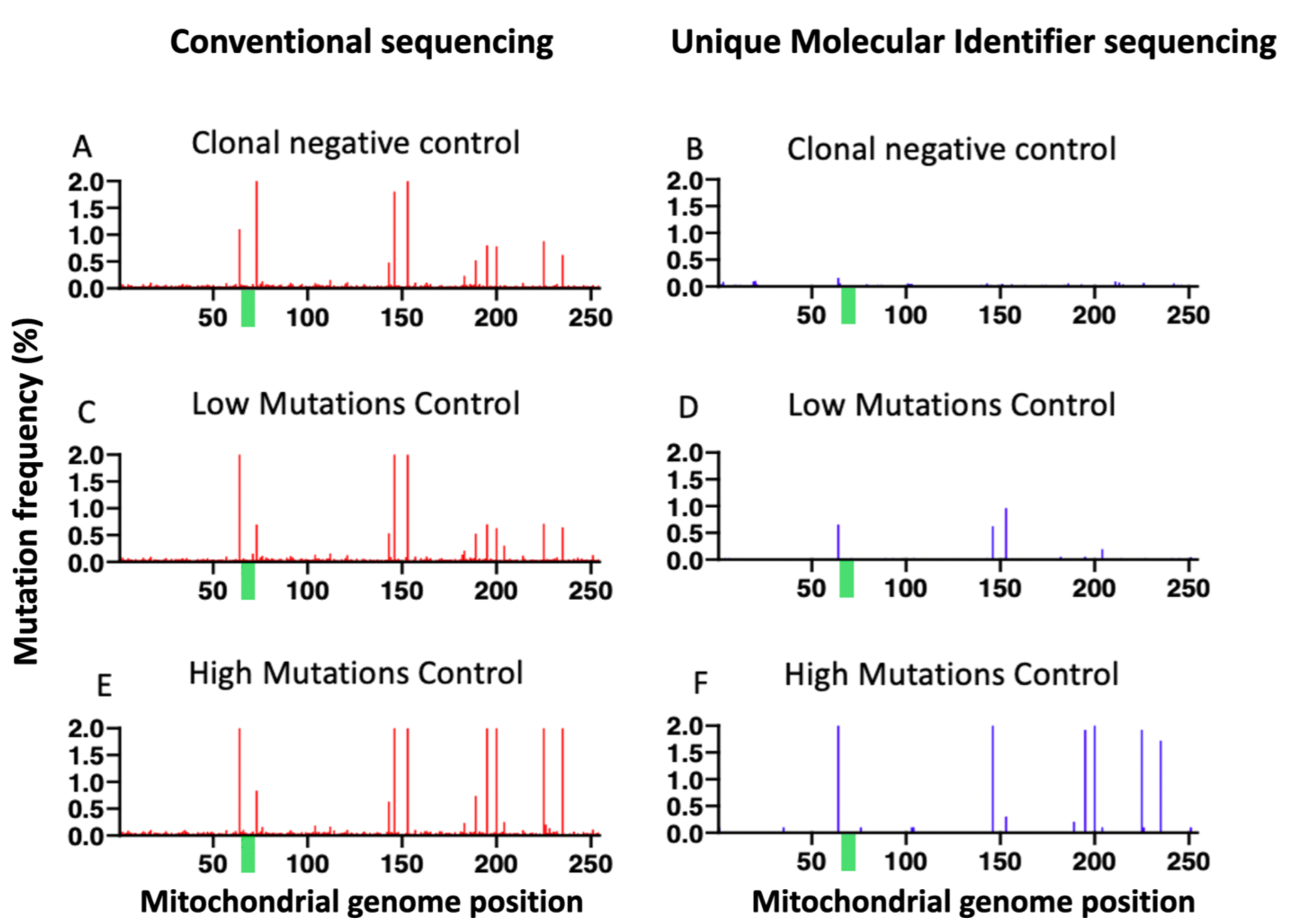
Mutation profile across the mCR using conventional NGS (A,C,E) compared to NGS with UMI applied (B,D,F). In a library prepared from a cloned plasmid, conventional sequencing (A) yielded 9 mutated positions with at least 0.5% prevalence, all of which are identified as artefacts and removed by UMI method (B). A similar mutation pattern is seen among other specimens (C,E), some of which is determined to be artefactual and subsequently removed with UMI sequencing (D,F). A homopolymer region of 6 Gs is indicated by the green bar. This is flanked on both sides by high prevalence mutations in all non-UMI applied samples. After UMI based sequencing, these mutations are removed in a clonal library, and only upstream mutations remain among the two pictured specimens.

Sequencing platforms such as Illumina are known to introduce sequencing errors associated with specific motifs^27^. One such region here is a 6bp G stretch (Figure 2, green bar) which may explain the mutations observed at flanking positions 64 and 73 (Figure 2A,C,E)^27^. Again, applying UMI-filtering successfully removed these in the cloned negative control (Figure 2B). The mutations remaining at position 64 and 73 in the high and low mutation controls post-UMI filtering (Figure 2D,F) may be represent Pol ɣ errors as the enzyme is more error-prone in homopolymer regions^28^. Variants detected in this region may therefore represent a combination of biological mutations and sequencing artefacts, and the UMI method presented here is capable of distinguishing between the two. UMI tagging allows reconstruction of a consensus sequence of all mtDNA molecule originating from a given individual specimen. This consensus can then be compared with individual mtDNA molecules represented by the UMI families from that same specimen, to identify mutations. This approach eliminates the need for sequence alignment and comparison to an arbitrary reference genome, and therefore allows the distinction between true mutations and polymorphisms which of course will differ between individuals, based on their maternal ancestry. This is an important advantage given that some ancestries are poorly represented in databases such as GenBank.

### Strengths and Limitations

The method presented here is designed to examine solely a targeted 264bp non-coding region within the mCR, where selective pressure is relatively low, and mutations and polymorphisms rates are prevalent. This method will therefore not inform about mutations in other regions of the mtDNA, including coding regions. The segment of interest was chosen for its very low likelihood of being deleted given its position within the regulatory region of mCR; deletion of this region would remove elements necessary for mtDNA replication. Another factor for choosing this segment was the rarity of previously reported mutations or polymorphisms within the region hybridizing with our assay primers. Indeed, having assaying approximately 3900 distinct individuals, we have never encountered a specimen for which the assay did not generate an amplicon. However, since the area of interest partially spans a hypervariable region, mutations in the primer regions are possible and if present, could impair UMI tagging, or alter the efficiency of the PCR stages, thus affect the detection of these mutations. It is also possible that polymerase errors may be introduced during the initial single amplification cycle used to tag mtDNA molecules. Such event would be rare and potentially difficult to distinguish from a true mutation, although as the mutation would be expected to appear in approximately 50% of reads, it would likely be rejected.

### Experimental design

#### Specimen requirements

Our protocol starts with total DNA extracted from fresh or frozen specimens; a wide range of tissue and DNA quality is acceptable, including fragmented or partially degraded DNA. As mtDNA copy number varies between tissues and specimen types, the starting quantity of tissue or cell number required for this method varies. We recommend using 3.5×10^6^ copies of mtDNA/μL for the initial tagging. Some types of specimens may present a challenge in attaining such mtDNA concentration. A lower mtDNA concentration is acceptable but may reduce the success of tagging and amplification stages; we have routinely obtained acceptable amplification of mtDNA with as low as 1.0 x10^5^ copies of mtDNA/μL when applying protocol adjustments (see notes in Box 1). For certain tissues or cell lines that have a high mtDNA copy number per cell, approximately 20ng of total DNA will provide sufficient mtDNA to carry out the assay but as little as 7ng of total DNA may suffice with some protocol adjustments (see notes in Box 1). For tissues with a lower mtDNA content, the recommended amount of total DNA is approximately 70ng, though as little as 25ng could be used with protocol adjustments.

In this protocol, two fragments, one in mCR and another within albumin (Alb) are amplified concurrently using monochrome multiplex qPCR with SYBR green fluorescence dye. This quantifies both mtDNA and nuclear DNA copy number, to determine mtDNA copies per nuclear genome, as described by Hsieh et al., 2018^29^. Since mtDNA copy number can vary between quantification techniques, we recommend using the aforementioned mtDNA quantification method for this protocol, although alternative methods can certainly be used. Though a lower amount of starting mtDNA may be acceptable, this would likely decrease the number of individually tagged mtDNA molecules, and potentially the total number of amplicons sequenced. This could in turn reduce the sequence coverage after raw reads are sorted into their sequence family (see Table 1 for term definitions). If a specimen were to harbor a variant within the primer annealing region, it may fail to amplify during the initial mtDNA quantification stage, or at the UMI labelling stage. The primers contain a few degenerate bases within the mtDNA complementary region to prevent the likelihood of this event. To date, in our hand, no specimen has ever failed to amplify.

#### Amplicon Tagging

A single PCR cycle takes place in the presence of 72bp degenerate primers. In addition to their complementary mtDNA sequence and their multiplex identifier (MID, see Table 1 for term definitions) to label each specimen, these elongation primers contain a UMI consisting of 13 random and 4 non-random positions. This generates 6.7 x 10^7^ individual unique barcodes that tag the mitochondrial genome starting at position M16559 (NC_012920.1), so that each DNA molecule can be uniquely identified with essentially no overlap in tagging sequences. The degenerate region of the primer contains two sets of 2bp anchor bases that facilitate alignment of the tag. From this point on, any errors introduced through PCR and/or sequencing can be identified during downstream bioinformatics analysis. The highest fidelity PCR polymerase available is used for this step, to minimize the potential of introducing a mutation during the initial amplification. Following this one round elongation tagging step, AMPure XP beads are used to remove any residual unincorporated primers and large nuclear DNA fragments. In this stage, library preparation negative controls are also prepared. These include one no-enzyme, one no-primer, and one no-DNA reaction. The maximum copy number observed in either the three negative controls is then subtracted from the specimen copy number, to determine subsequent dilution steps. After clean-up of the labelled products, a second PCR step amplifies and adds the multiplexing indices, needed for Illumina sequencing to each labelled template.

#### UMI-tagged templates amplification for Sequencing

Following quantification, each UMI tagged amplicon is diluted to the desired concentration and then amplified for Illumina MiSeq sequencing. The number of labeled template copies used to this step will determine both the family size (see Table 1 for term definitions) and the number of resulting read families, thereby determining the overall sequence coverage of each DNA extract. A lower concentration of templates at this step tends to result in a lower number of UMI families, and these families contain a larger number of reads. Conversely, at very high concentrations, a substantial proportion of reads belong to families that count fewer than 5 reads hence are discarded (Figure 3).

**Figure 3:**
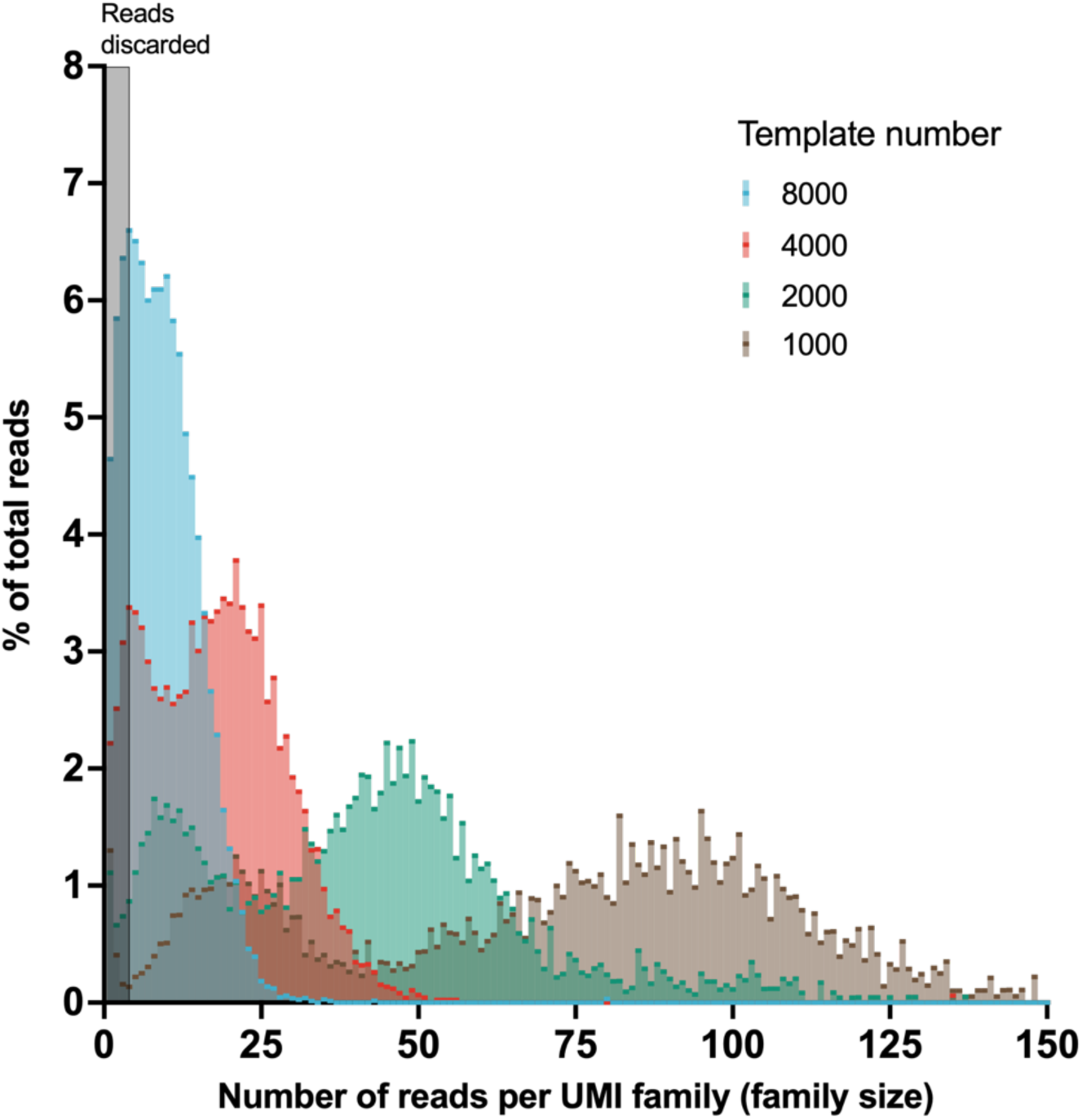
Initial template quantity determines family size. Sequencing 1000 UMI-tagged templates results in allocation of a large percentage of reads being assigned to needlessly large families. Increasing the number of UMI-tagged templates results in a greater proportion of reads allocated to smaller families. However, sequencing 8000 UMI-tagged templates leads to a large proportion of reads being allocated to families that contain fewer than 5 reads, hence are not usable.

Families containing 5 to 10 reads are identified as the optimal family size, contributing to an increased effective depth of coverage. Large families, especially those with over 25 reads per UNI family, indicate a suboptimal distribution of reads which decreases the number of usable UMI families, resulting in a reduction of coverage. Allocating excess reads to smaller families is beneficial, as it enhances the number of families surpassing the family size threshold. When a lower number of UMI-tagged templates are amplified by PCR (i.e. 1000 and 2000), we observe that a substantial proportion of families are assigned between 50 to 100 reads (Figure 3). As the number of templates increases, the distribution of reads shifts towards smaller families (Figure 3), although this can eventually increase the proportion of families containing fewer than the recommended threshold, making them unusable. We advocate for at least 5 reads per family, a rationale that will be discussed in the following sections. If using our specified analysis parameters, our recommendation is to add 4000 UMI-tagged templates at the PCR stage, as this yields the largest number of families with a sufficient size (Figure 3). This template number may be adjusted depending on the desired MiSeq platform and the number of specimens per sequencing run.

### Bioinformatic Analysis Pipeline

To analyze raw sequencing data, we have developed a pipeline (https://github.com/BoxeRZL/UMI-DloopSeq) built primarily with the python package Biopython. Analysis is conducted through five steps (Figure 4). The first step involves trimming raw FASTQ sequences and combining trimmed reads, which also provides a quality control summary. The second step consists of sorting and grouping surviving reads, first based on MID, then based on the UMI barcode. In this step, ungrouped sequences or sequences without a recognizable barcode are excluded. The barcode sequences are recognizable by the presence of anchor bases interspersed among the degenerate regions as such (N)^4^CA(N)^4^GT(N)^5^ (Table 2). The absence of any of these indicates improper barcode labelling. Additionally, this step includes an initial alignment of reads within each family, to form a family consensus (see Table 1 for term definitions) sequence, and a subsequent re-alignment to identify any ambiguous positions where the base call is not definitive. The third step summarizes the distribution of the family size across specimens. We recommend a threshold of at least 400 UMI families; fewer families may introduce bias, as a small fraction of initial templates may be overrepresented. The fourth step calls low frequency mutations from the alignment of family consensuses. A mutation is confirmed if it is generated from a UMI family with at least 5 reads, and if it is present in at least 75% of the reads within that family. The final step estimates the overall mutation burden, which quantifies the total number of mutations per 10,000 bp in the sequenced region.

**Figure 4:**
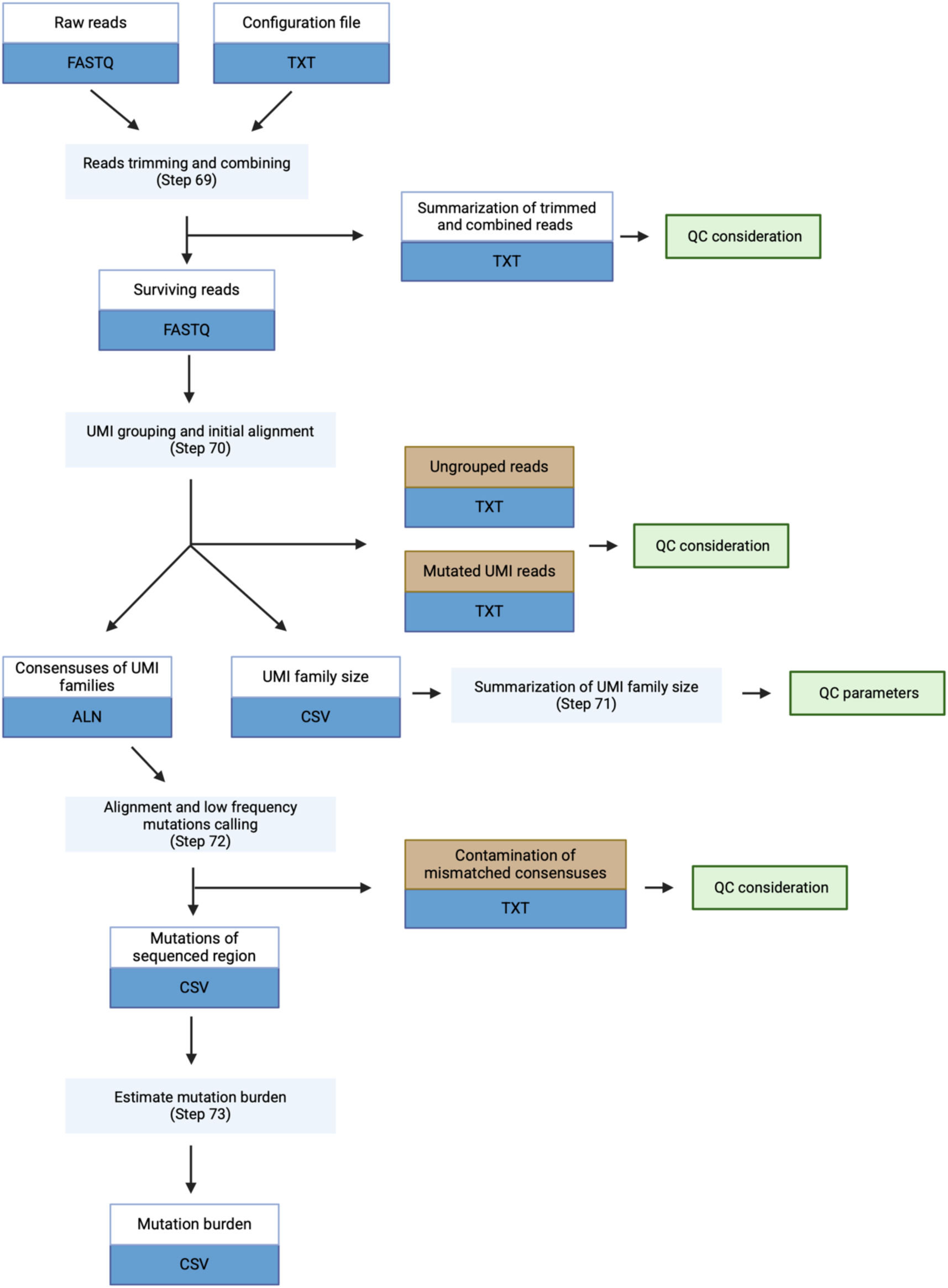
Flowchart of the bioinformatics pipeline. The pipeline includes five main steps. The light blue rectangles show data operations. The white & blue rectangles show explanations and types of data obtained. The brown & blue rectangles show reads and/or consensuses excluded from the pipeline. The green rectangles show consideration and/or parameters processed for quality control (QC).

**Table 2:**
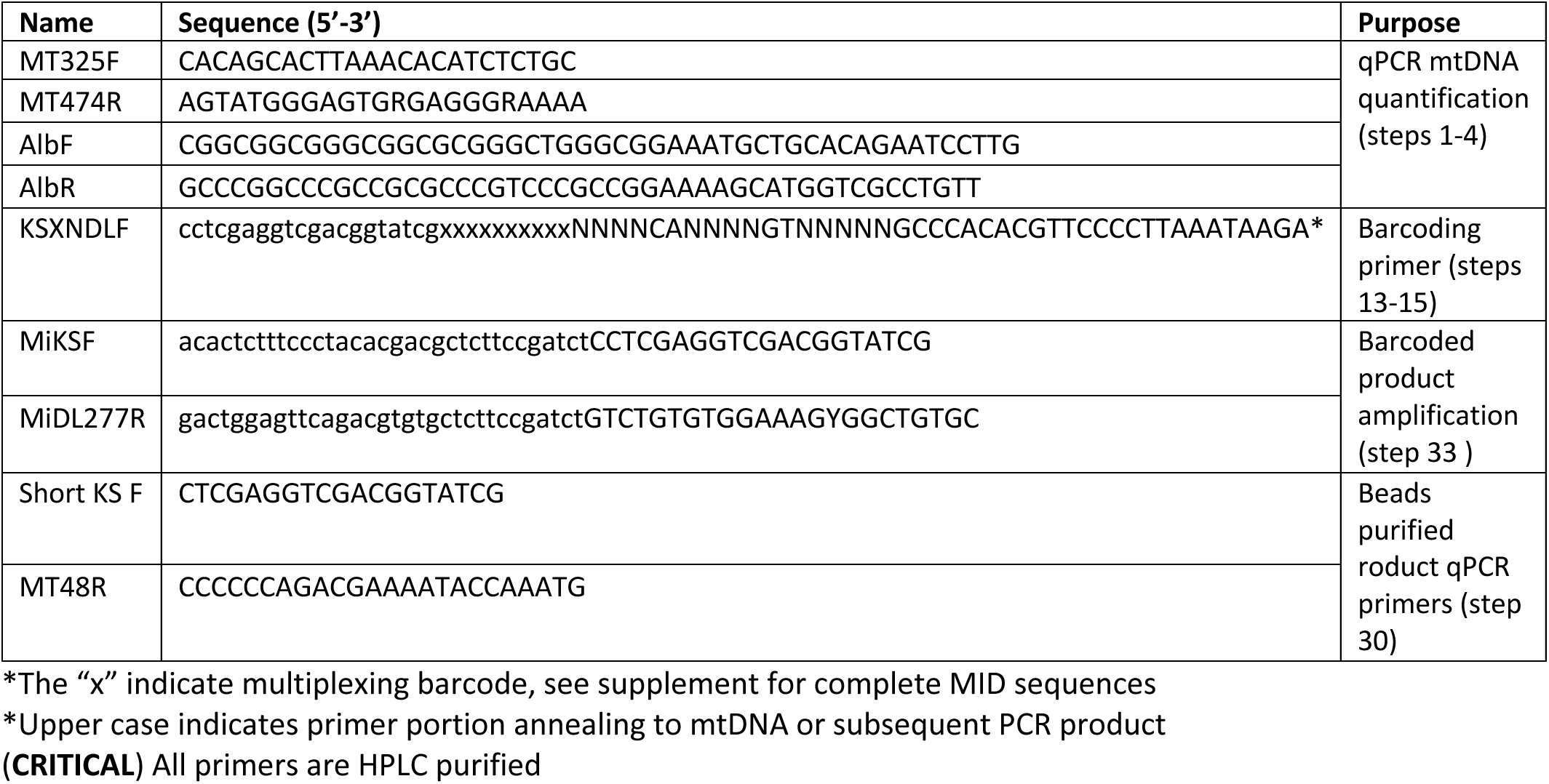
List of primer sequences, and associated protocol steps.

### Read trimming and alignment

Our analysis pipeline employs Trimmomatic^30^ and FLASH^31^ for the initial preprocessing of raw sequences, ensuring sufficient read quality and combining the paired 2X300 MiSeq reads. The processed reads are sorted based on the MID sequence, then based on the UMI sequence, effectively grouping them into corresponding read families. Utilizing the MiSeq 2X300 platform, with an allocation of 300 specimens per flow cell, a loss of approximately 33% of reads after trimming and combining of paired reads is expected. For poor quality sequencing data, the stringency of reads trimming and combining threshold may be decreased. However, this adjustment may increase the susceptibility of the resulting analysis to errors, particularly if the sequence quality is low. For a 300-specimen run using 4000 UMI-tagged templates, our post-analysis results yield an average of approximately 1000 unique barcodes represented with a sufficient family size. This allows for a minimum resolution of mutations present at 1 in 1000, or ∼0.1%. To increase the coverage of specimen consensus (see Table 1 for term definitions) sequences, alterations can be made, such as decreasing the number of specimens in a single sequencing run and/or increasing the number of barcoded templates sequenced (Figure 3).

#### Low frequency mutations calling

To enhance the precision of mutation calling at individual positions and mitigate background noise in family consensus sequences, we conducted a comprehensive evaluation of analysis parameters designed to minimize artefacts arising from PCR and/or sequencing errors. We investigated the effects of family size and intra-family agreement, the latter reflecting the frequency of base occurrence required to generate a consensus sequence within a family. Our results demonstrate that a maximum intra-family agreement threshold effectively eliminates erroneous data in the clonal library while reducing the sensitivity of mutation detection in internal control (Figure 5A and C). Specifically, setting a family size threshold of 5, combined with a 75% intra-family agreement, substantially reduces the presence of false positive mutated bases and optimizes the number of usable UMI families (Figure 5B). Further comparative analysis showed that variations in intra-family agreement thresholds have a more marked impact on mutation calling outcomes and the coefficient of variation (CV) in internal control than adjustments in family size thresholds (Figures 5C, D, and E).

**Figure 5:**
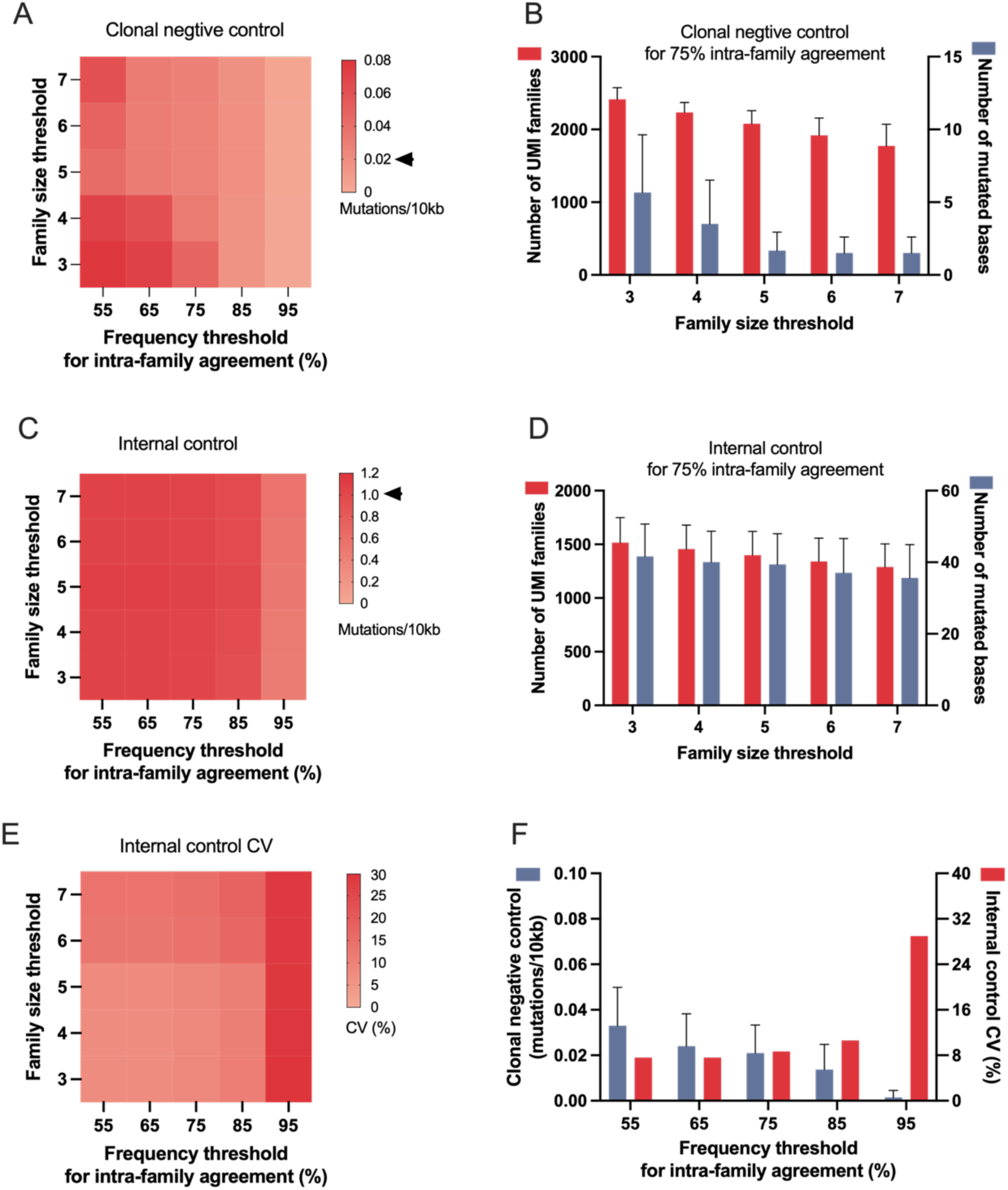
OpKmizaKon of assay parameters enhances the detecKon of low frequency mutaKons. (A) Heatmap showing that increasing thresholds of family size and frequency of base occurrence accepted for generating a family consensus sequence reduce false positivity of mutation calling in the clonal library. The black arrow indicates the expected result of false positivity with the optimized combination of family size and intra-family agreement thresholds. (B) With frequency for intra-family agreement set to 75%, a family size threshold of 5 effectively excludes most erroneous mutations while retaining a sufficient number of usable UMI families. (C) A slight reduction in mutation burden in internal control is associated with increased family size threshold, given a fixed intra-family agreement frequency. Notably, seqng the intra-family agreement frequency at 95% results in lower mutation detection in internal control. The black arrow indicates the expected mutation burden of internal control with the optimized combination of family size and intra-family agreement thresholds. (D) With frequency for intra-family agreement set to 75%, the number of usable UMI families and detected mutated bases gradually decreased as the threshold of accepted minimum family size increases. (E) Heatmap showing that the increase in the coefficient of variation (CV) for mutation detection in internal control, associated with rising thresholds of family size and intra-family agreement frequency. (F) With family size set to 5, an intra-family agreement frequency threshold of 75% optimizes the trade-off between minimizing background noise and reducing the CV of mutation detection in internal control.

To effectively optimize these analysis threshold parameters, it is essential to consider the balancing effects of threshold parameters. Increasing the family size threshold improves the accuracy of generating family consensus sequences by elevating the minimum required number of reads within a family. However, with a lower intra-family agreement threshold of 55%, the overall mutation burden from false positives increases as the family size threshold is raised from 5 to 7, due to a concurrent reduction in the number of usable UMI families (Figure 5A). Conversely, raising the intra-family agreement threshold enhances the precision of base identification at each position within family consensus sequences, which reduces false positivity across the entire sequencing region of interest. However, setting the intra-family agreement to 95% dramatically restricts mutation detection sensitivity, and increases the CV in internal control, potentially leading to the exclusion of many true mutations (Figures 5C and E). It is also important to note that if using smaller families, the threshold for the intra-family agreement percentage should be set keeping in mind that a single mutated read can disproportionately affect the results.

In conclusion, to strike an optimal balance between minimizing false positives and ensuring accuracy in mutation detection, we found that, under the conditions of our assay, maintaining a family size threshold of five reads or more per family and a 75% or more intra-family agreement threshold is optimal (Figure 5F). The black arrows highlight the expected results of false positivity in the clonal library (Figure 5A) and mutation burden in internal control (Figure 5C).

### Quality control considerations

#### Misalignment

Misalignment of sequences has been a persistent challenge in this field of research, given that any misalignment could greatly inflate the identification of low frequency mutations, hence the estimation of overall mutation burden. To mitigate this, our methodology includes a re-alignment step, after construction of the family consensus. In this step, if a specific base at a given position cannot be definitively identified by the read alignment within the family, it will be temporarily substituted with an N. To minimize erroneous classification of these ambiguous bases as true mutations, Ns are reverted to reflect the specimen consensus sequence. This conservative approach may slightly underestimate the mutation burden, but it minimizes variability that could potentially be introduced by weak agreement within families due to either poor sequencing data quality, or contamination introduced at any stage.

#### Contaminant filtering

In order to minimize “contamination” by non-mtDNA sequences that shares homology with human mtDNA, we implemented an identity threshold for the pairwise alignment. Our analysis revealed that, on average, there was < 1 non-mtDNA sequence per DNA specimen, or approximately per 2000 family consensuses. Utilizing BLAST, we identified that these sequences usually were from bacterial DNA, likely a result of specimen contamination by bacteria that share sequence homology with human mtDNA, and are amplified early in the library preparation process. To exclude bacterial sequences, family consensuses with less than 80% identity to a specimen’s mtDNA consensus were removed.

In the analysis of whole blood DNA, we also identified a nuclear mitochondrial DNA (NUMT) region of chromosome 17 that shares 89% identity with the corresponding region of the mCR. In tissues where NUMTs were detected, or in those with a lower mtDNA content, we raised the contamination threshold to over 95% sequence identity, to ensure the exclusion of these sequences. Finally, to mitigate the effect of potential index hopping events^25,26^, if a read within a UMI family counts more than 15 positions (15/264) showing a mutation, that read is excluded.

The presence of contaminants is a critical factor in the consideration of quality control; a high number of these contaminating sequences may indicate poor conditions during library preparation.

#### Expertise recommended to implement the protocol

This protocol outlines steps for amplicon library preparation and post-sequencing bioinformatics analysis. Sequencing of the amplicons itself should be performed by individuals highly experienced with next generation sequencing, such as a core facility specialized in sequencing. For example, we used the Sequencing and Bioinformatics Consortium at the University of British Columbia, Vancouver, BC, Canada.

#### Reagents

##### DNA quantification

- LightCycler 480 SYBR Green I Master (Roche, cat.no. 04887352001)
- TOPO-D-loop Plasmid DNA Standard dilutions (from 3.54e7 copies/2μL to 3.54e1 copies/2μL) prepared as described in Hsieh et al., 2018
- MtDNA qPCR internal control prepared as described in Hsieh et al., 2018
- Dloop (mCR) Primer mix: MT325F + MT474R, 10μM each (Table 2)
- Albumin Primer mix: AlbF + AlbR, 10μM each (Table 2)

##### Targeted DNA region extension

- KSXNDLF forward primer, 25μM (Table 2)
- PFuUltra II Polymerase (Agilent, cat.no.600672 / 200rxs; 600674 / 400rxs)
- 10X PfuUltra II Reaction Buffer (Agilent, cat.no.600670)
- dNTP (25mM) (Invitrogen, cat.no.55084)

##### AMPure beads purification

- Agencourt AMPure XP beads, 5 mL (Beckman Coulter, cat.no.A63880)

##### DNA amplification

- KS Primer mix (Short KSF + MT48R), 10μM (Table 2)
- Mi Primer mix (MiKSF + MiDL277R), 25μM (Table 2)
- PFuUltra II Polymerase (Agilent, cat.no.600672 / 200rxs; 600674 / 400rxs)
- 10X PfuUltra II Reaction Buffer (Agilent, cat.no.600670)
- dNTP (25mM) (Invitrogen, cat.no.55084)

##### Gel electrophoresis

- GelRed Nucleic Acid Stain (BIOTIUM, cat.no. 41003)
- UltraPure Agarose (Invitrogen, cat.no. 16500-500)
- 1X Tris-acetate-EDTA (TAE) Buffer (Referred to reagent set up section)
- 10X loading dye (Invitrogen, cat.no.10816015)

##### Gel extraction

- QIAquick Gel extraction kit (QIAGEN, cat.no.28704)
- 2-propanol (Sigma, cat.no.I9516) (CAUTION: 2-propanol is flammable, so avoid heat, spark, and open flame when using it.)

##### General reagents

- Buffer AE (QIAGEN, cat.no.19077)
- DNase, RNase-free UltraPure Water (Invitrogen, cat.no.10977)
- TRIS-HCl pH 8.0 buffer, 10 mM (Sigma, cat.no.T3038)
- Buffer TE (contains 10 mM Tris-Ace, 1 mM EDTA, pH 8.0) (Thermo Fisher Scientific, cat.no.AM9849)
- DNAZap™ PCR DNA Degradation Solutions (Ambion, cat.no.AM9890)
- 95% & 70% ethyl alcohol (Commercial Alcohols, cat.no.P016EA95) (CAUTION: Ethyl alcohol is flammable, so avoid heat, spark, and open flame when using it. Ethyl alcohol is also irritating to the skin and eyes)

#### Materials

##### DNA quantification

- LightCycler 480 multiwell plate, white (Roche, cat.no. 04729692001)
- LightCycler 480 Sealing Foil (Roche, cat.no.04729757001)

##### General materials

- 0.2mL PCR tubes, DNAse/RNAse-free (Axygen, cat.no. 321-02-051)
- 1.5 mL screw-cap tubes (Sarstedt, cat.72.692.405 or 72.692.415)
- 1.7mL ultraclear snap-cap microtubes, DNAse/RNAse-free (Progene, cat.no.87-B175-C)
- 2.0 mL screw-cap tubes (Sarstedt, cat.no.72.693.465)
- Falcon tube, 50 mL (Avantor, cat.no.525-1074)
- Polystyrene serological pipettes, 25 mL, 5 mL, 2 mL (FroggaBio, cat.no.SP25, cat.no.SP5, cat.no.SP2)
- Low retention pipette filter tips: P10, P20, P100, P200, and P1000 (FroggaBio, cat.no.FT10, cat.no.FT20, cat.no.FT100, cat.no.FT200, cat.no.FT1000)
- Thermowell Sealing Tape (Corning, cat.no. 6569)
- Powder-free nitrile gloves (SensiCare, cat.no.MDS6803)
- Paper towels (generic)
- Lint-free Kimwipes (Kimberly-Clark, cat.no.34256)
- Plastic Wrap (Glad, Cling wrap)
- Absorbent Underpad (Fisher Scientific, cat.no.14-206-65)
- Scalpel blades (size 22) (Integra, cat.no.4-122)
- Weighing boat (Fisher Scientific, cat.no.2-202A)
- Protective sleeves, Tyvek (VWR, cat.no.414004-420)
- Masking Tape (generic)
- Ice

#### Equipment

##### DNA quantification

- LightCycler 480 II (Roche, cat.no.29579)

##### Targeted DNA region extension

- MyCycler, Thermal Cycler (Bio-Rad, cat.no.580BR7544)

##### AMPure beads purification

- DynaMag™-2 Magnet (Thermo Fisher Scientific, cat.no. 12321D)

##### DNA amplification

- MyCycler, Thermal Cycler (Bio-Rad, cat.no.580BR7544)

##### Gel electrophoresis

- 250mL Erlenmeyer flask (Pyrex, cat.no.4980)
- Gel casting tray (Bio-Rad, cat.no.1704426)
- Gel Electrophoresis power supply (E-C Apparatus, cat.no.EC105)
- Gel Electrophoresis apparatus (Bio-Rad, cat.no.258BR013827)

##### Gel extraction

- UV transilluminator (302nm) (UVP, cat.no.042905-002)
- Balance (Sartorius, cat.no.TE412)
- Safety goggles, with UV filter (North, cat.no.UV50LG/N)
- Heat block (temperature range of 0 C to 100 C) (VWR, cat.no.13259-032)

##### General equipment

- Eppendorf centrifuge 5810R (Eppendorf, cat.no.0031888)
- Eppendorf centrifuge 5415D (Eppendorf, cat.no.0060958)
- VORTEX-Genie 2 (Scientific Industries, cat.no.G-560)
- Pipetman Single Channel Classic Pipettes: P2, P10, P20, P100, P200, and P1000 (Gilson, cat.no.Model P2, cat.no.Model P10, cat.no.Model P20, cat.no.Model P100, cat.no.Model P200, and cat.no.Model P1000)
- VWR Galaxy Mini centrifuge (VWR, cat.no.SN07021238)
- Pipette Filler (Thermo Scientific, cat.no285365)
- Seal applicator (generic)
- -20 °C freezer (any brand)
- -80 °C freezer (any brand)

#### Data analysis

##### DNA quantification: software

- LC480 Conversion 2.0 (https://medischebiologie.nl/files/?main=files&sub=0)
- LightCycler software (https://lifescience.roche.com/global/en/products/product-category/lightcycler.html#software)
- LinRegPCR software 2012.1 (https://medischebiologie.nl/files/?main=files&sub=0)

##### Bioinformatics analysis: software

- Python 3.7 or later (https://www.python.org/downloads/)
- Python package Biopython 1.75 or later(https://biopython.org)
- Trimmomatic 0.36 or later(http://www.usadellab.org/cms/?page=trimmomatic)
- Mafft 1.6 or later (https://mafft.cbrc.jp/alignment/software/source.html)
- Flash 1.2.11 or later (https://ccb.jhu.edu/software/FLASH/)
- Slurm 23.11 or later (https://slurm.schedmd.com/documentation.html)

##### Bioinformatics analysis: pipeline

- UMI-DloopSeq (https://github.com/BoxeRZL/UMI-DloopSeq)

Reagent setup

50X TAE buffer (1 L)

- 242 g UltrPure Tris (Invitrogen, cat.no.15504-020)
- 57.1 mL Acetic Acid, Glacial (Fisher Scientific, cat.no.A38212)
- 100 mL Ethylenediaminetetraacetic acid disodium salt solution (EDTA), 0.5M (Sigma, cat.no.E7889)

1X TAE buffer

Take 100 ml of the 50X TAE and add 4900 mL water

#### Procedure

##### Quantification of mtDNA copy number

1. Thaw, vortex and give a quick spin to qPCR primers (MT325F + MT474R, AlbF + AlbR), Roche SYBR green II master, and DNA to be assayed
2. Dilute all DNA extracts 1:10 (this may not be required but is recommended unless the DNA concentration is very low)
3. Set up 96 well plate layout, include 7 standards (preparation in **Box 1**) in single, one negative control in single, two internal controls each in duplicate, and each DNA extract to be assayed, in duplicate (the number of wells used in the plate will be 2X+12 where X is the number of specimens being assayed; example of plate layout shown in supplement F1)
4. Prepare the master mix according to the following Table, including all components except the DNA, and multiplying each volume by the number of wells that will be used plus a few (usually 96 + 4 = 100) to ensure sufficient volume remaining by well 96. Pipette the master mix up and down gently to mix then spin down and load 8 μL into each well of the 96 well plate.

| Component | Assay Final Concentration | Volume (µL) |
| --- | --- | --- |
| LightCycler 480 SYBR Green I Master | 1X | 5 |
| PCR-grade Water | - | 2 |
| Primer Mix (MT325F + MT474R, AlbF + AlbR) | 1 µM | 1 |
| DNA | Variable | 2 |

**(CRITICAL STEP) hold the multiwell plate by its sides during handling**

5. Deliver 2 μL of DNA to be assayed in each respective well
6. Deliver standards, internal controls, and negative control to their respective well
7. Seal the white multiwell plate with LightCycler 480 sealing foil, and press with seal applicator
8. Centrifuge for 2 min at 1500xg (Eppendorf 5810R), repeat if there are bubbles in the wells
9. Insert the plate into the LightCycler 480 machine. The program shown in the table below should have been pre-programmed. Begin the run

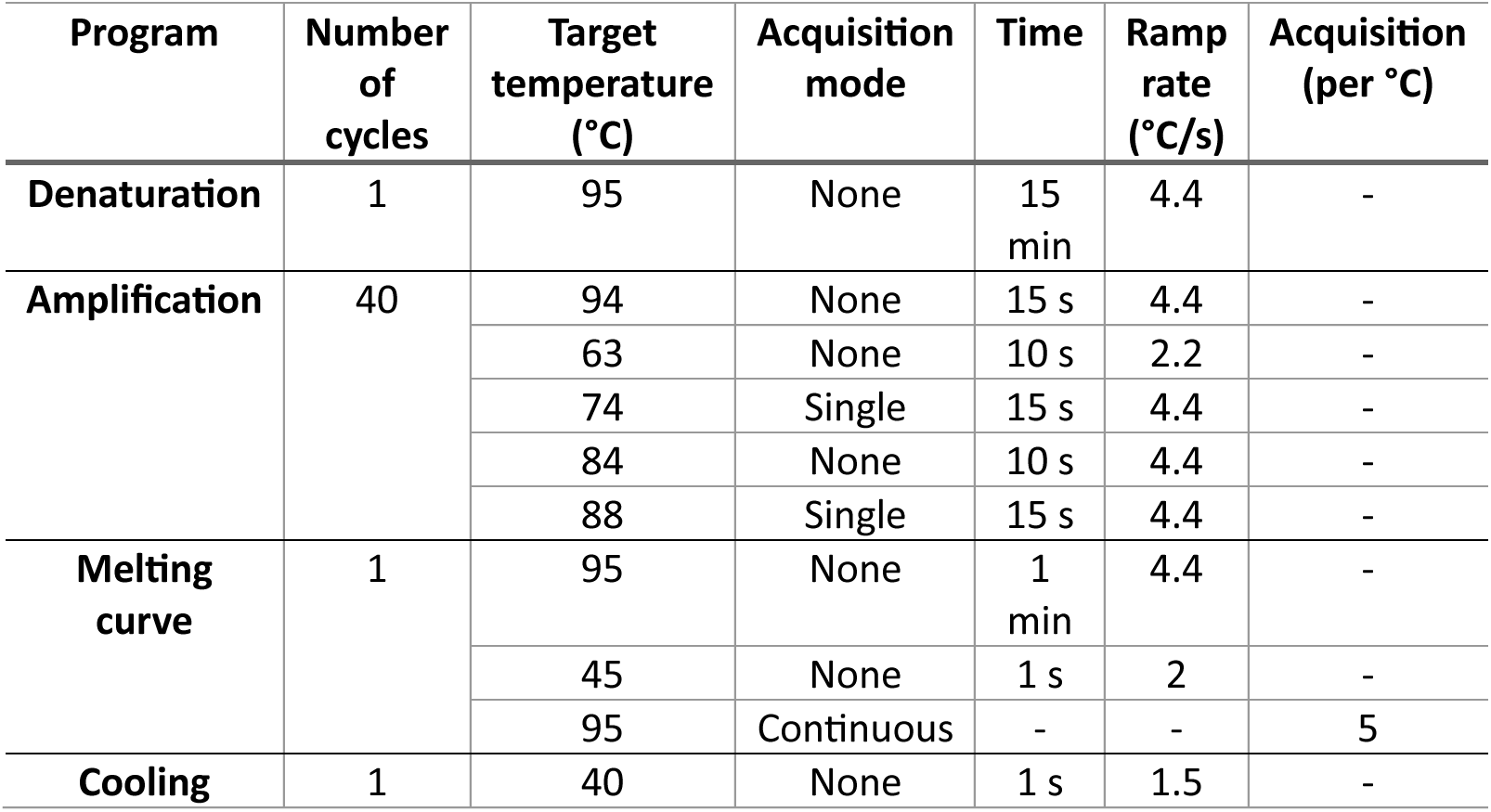

| Program | Number of cycles | Target temperature (°C) | Acquisition mode | Time | Ramp rate (°C/s) | Acquisition (per °C) |
| --- | --- | --- | --- | --- | --- | --- |
| **Denaturation** | 1 | 95 | None | 15 min | 4.4 | - |
| **Amplification** | 40 | 94 | None | 15 s | 4.4 | - |
|  |  | 63 | None | 10 s | 2.2 | - |
|  |  | 74 | Single | 15 s | 4.4 | - |
|  |  | 84 | None | 10 s | 4.4 | - |
|  |  | 88 | Single | 15 s | 4.4 | - |
| **Melting curve** | 1 | 95 | None | 1 min | 4.4 | - |
|  |  | 45 | None | 1 s | 2 | - |
|  |  | 95 | Continuous | - | - | 5 |
| **Cooling** | 1 | 40 | None | 1 s | 1.5 | - |

The outcome data of the LightCycler 480 software contains seven columns, including sample position, sample name, program, segment, cycle, time, and temperature. Signals of the mCR (74 °C) and Alb (88 °C) fragments are delineated with markers of the program and segment indicated by the LightCycler 480 software. Utilizing the thermal cycling described in step 8, data of mCR fragment are shown in program 2 and segment 3, whereas data of Alb fragment are shown in program 2 and segment 5. The separated mCR fragment data are converted to grid format by LC480 Conversion software. The calculation of mtDNA copy number is performed based on converted data with the baseline correction by LinRegPCR software as described by Hsieh et al., 2018^29^.

10. Based on the mtDNA concentration determined, dilute each DNA in AE buffer to 1,400,000 mtDNA copies/μL. You will need 2.5μL of the diluted DNA for subsequent steps **(CRITICAL STEP) for specimens with lower concentrations, do not dilute, follow additional instructions in Box 1)**

**(PAUSE POINT) samples may be kept at -20 °C indefinitely (for long periods store at -80°C to prevent evaporation)**

##### Extension of targeted region in mtDNA tagging with MID and UMI

11. Randomize all DNA samples and assign a multiplexing primer, including MID and UMI, and lane number. We recommend assigning 12 DNA samples per lane (this can be increased up to 43, see supplement for extended list of multiplexing primers). The number of lanes in a single MiSeq run will depend on the desired sequencing depth, we recommend sequencing 25 lanes of 12 multiplexed samples (or 7 lanes of 43 multiplexed samples) for 300 specimens per MiSeq v2 2X300bp run.

**(CRITICAL STEP) From this point forward, any cross contamination between lanes will not be distinguishable from real mutations. Clean all benches and pipettes with DNAzap for 5 min, wipe with paper towel then rinse thoroughly with 70% ethyl alcohol to remove residual DNA and DNAzap. Alternatively, benches may be cleaned with bleach for 5 min and rinsed with ethyl alcohol as above or use disposable bench protectors and change between each use.**

**(CRITICAL STEP) Step 12 is presented for the preparation of a single specimen, calculate the master mix volumes according to the number of DNA samples to be multiplexed per lane. Include 3 controls: no enzyme, no primer, and no DNA negative control.**

12. Thaw all reagents and mix components while on ice according to the table below.

**(CRITICAL STEP) PfuUltra II polymerase must be kept at -20°C, make sure the master mix is cold before adding the enzyme, and keep tubes on ice at all times.**

| Reagent | Final Concentration | Volume (μL) |
| --- | --- | --- |
| UltraPure water | - | 17.25 |
| 10X PfuUltra II Reaction Buffer | 1X | 2.50 |
| dNTP (25 mM) | 0.25 mM | 0.25 |
| PfuUltra II polymerase | 1U | 0.50 |

13. Aliquot 20.5μL of master mix into labeled 0.2mL PCR tubes, add 2.5μL of diluted DNA from step 10 to the master mix and 2μL of the respective multiplexing primers diluted to 6.25μM. See supplement for a list of multiplexing primers with MID sequences.
14. Vortex briefly on low speed, or pipette up and down gently to mix without denaturing the enzyme.
15. Place tubes in thermocycler and run the PCR program:

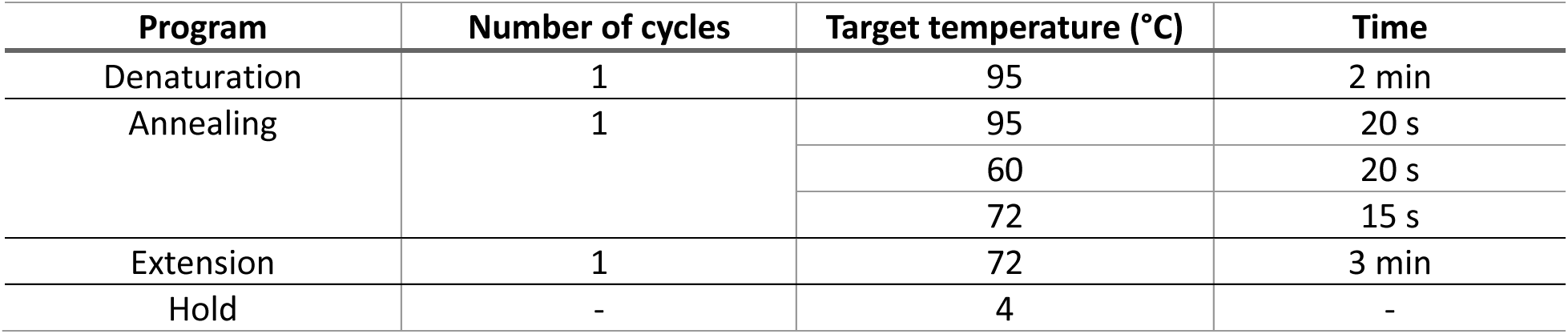

**(PAUSE POINT) samples may be stored at -20 °C for several weeks (for long periods store at -80°C to prevent evaporation)**

##### AMPure beads purification

16. Label 1.7mL snap cap tubes, 12 for each sample and 3 for the no DNA, no primer, and no enzyme controls, then add 50μL UltraPure water to each **(CRITICAL STEP) use 96 wells plates when the number of samples is increased to 43**
17. Add 50μL AMPure beads to each tube
**18.** Transfer 25μL of the PCR product from step **15** into their respective tubes and pipet up and down 10 times to mix, this is preferable to vortexing as it is gentler on the DNA and minimizes risk of droplets **(CRITICAL STEP: Ensure you change tip for each sample)**
19. Incubate for 10 min at room temperature for maximum recovery
20. Briefly spin down to remove any droplets from the sides and cap, then place in DynaMag-2 magnet for 5 min. The solution should go from cloudy to clear with a distinct pellet containing the beads and bound DNA.
21. Remove and discard the supernatant without disturbing the pellet, avoid touching the rims or caps as this can be a source of contamination
22. Remove tubes from magnet rack and add 190μL of 70% **ethyl alcohol**, gently vortex for 5 s to resuspend. (**CRITICAL STEP) vortex an additional 5 s if pellet is not completely resuspended**
23. Spin down briefly to remove droplets from the sides and cap, then replace in DynaMag-2 magnet for 5 min
24. Do not remove tubes from magnet rack, with a P200 remove and discard supernatant without disturbing the pellet
25. Repeat steps 22-24, be sure to remove as much supernatant as possible without disturbing the pellet
26. Keeping the tubes in the magnet rack, open the caps to allow pellets to dry. When the pellet is dry it will turn from shiny to matte, if the pellet becomes overdried and cracks this will reduce yield. To speed up this step the tubes may also be placed in a heating block set to 37 °C to dry.
27. Add 30μL of buffer TE and vortex for 5 s
28. Briefly spin down and place tubes on magnet rack for 5 min
29. Transfer the supernatant into prelabeled 1.5mL screw cap tubes **(CRITICAL STEP) do not use snap cap tubes as droplets may fly out when opening the caps**

**(PAUSE POINT) tubes may be stored at -20 °C for several weeks (for long periods store at -80°C to prevent evaporation)**

##### Quantification and amplification of products purified by AMPure beads

30. Quantify AMPure purified product from step 29, prepare master mix components following steps 3-9 this time with the KS primer mix (Short KSF + MT48R). Include the no DNA, no primer, and no enzyme controls. The program shown in the table below should have been pre- programmed.

| Program | Number of cycles | Target temperature (°C) | Acquisition mode | Time | Ramp rate (°C/s) | Acquisition (per °C) |
| --- | --- | --- | --- | --- | --- | --- |
| Denaturation | 1 | 95 | None | 10 min | 4.4 | - |
| Amplification | 45 | 95 | None | 5 s | 4.4 | - |
|  |  | 60 | None | 10 s | 2.2 | - |
|  |  | 72 | Single | 5 s | 4.4 | - |
| Melting curve | 1 | 95 | None | 1 min | 4.4 | - |
|  |  | 45 | None | 1 s | 2 | - |
|  |  | 95 | Continuous | - | - | 5 |
| Cooling | 1 | 40 | None | 1 s | 1.5 | - |

31. Determine the background which will be the maximum copy number value among the no DNA, no primer, and no enzyme controls, then subtract that background from the copy number of each sample, to find the resulting copies/μL
32. Dilute the AMPure purified product to 4000 copies per reaction (or 400 copies/μL)

**(CRITICAL STEP) the dilution concentration may be adjusted depending on the desired depth of sequencing, a higher concentration of DNA will result in higher sequencing coverage depth.**

33. Prepare PCR reaction master mix in a clean area (this should be free of PCR products), and load 40μL into each 0.2mL PCR tube

| Component | Final Concentration | Volume per reaction |
| --- | --- | --- |
| UltraPure water | - | 31.5 |
| 10X PfuUltra II Reaction Buffer | 1X | 5.0 |
| dNTP (25 mM) | 0.25 mM | 0.5 |
| MiKSF (25 µM) | 0.5µM | 1.0 |
| MiDL277R (25 µM) | 0.5µM | 1.0 |
| PfuUltra II polymerase | 1U | 1.0 |

34. Load 10μL of the DNA from step 32 into their respective tubes, and pipette to mix thoroughly
35. Load tubes into a thermocycler with the following cycling program:

34. Load 10µL of the DNA from step 32 into their respective tubes, and pipette to mix thoroughly 35. Load tubes into a thermocycler with the following cycling program:
| Program | Number of cycles | Target temperature (°C) | Time |
| --- | --- | --- | --- |
| Denaturation | 1 | 95 | 2 min |
| Annealing | 35 | 95 | 20 s |
|  |  | 60 | 20 s |
|  |  | 72 | 15 s |
| Extension | 1 | 72 | 3 min |
| Hold | - | 4 | - |

**(PAUSE POINT) After the PCR, tubes with DNA may be stored at -20 °C until ready for next step, for storage longer than 2-3 days transfer to screw top tubes and store at -80°C to prevent evaporate Gel electrophoresis and extraction of PCR amplicons and sequencing**

**(CRITICAL STEP) ensure all gel electrophoresis apparatus and bench surfaces have been cleaned with DNAZap before starting and between each use**

36. Make a 2% gel by mixing 3.00g of UltraPure agarose with 150mL of 1X TAE in a 250mL Erlenmeyer flask
37. Cover the flask with wet paper towel to prevent evaporation and microwave for 3 min or until all the agarose is melted and solution is clear, checking and mixing every 30 s
38. Add 15μL of GelRed to melted agarose solution, swirl to mix
39. Place flask in dark area to allow it to cool slightly
40. Prepare gel casting tray by taping the sides, and obtain clear gel comb with 20 teeth, 7mm wide and 1.5mm deep
41. Once the molten gel in the flask has cooled so it is comfortable to the touch, pour it into the gel casting tray ensuring there are no bubbles
42. Leave gel to solidify 0.5-1h while protected from light
43. Thaw PCR products from step 35, vortex and spin down volume
44. For each set of 12 multiplexed samples pool 10μL from each tube, for a volume of 120μL
45. Remove tape and comb from solidified gel, place it in electrophoresis chamber, and add 1X TAE until gel is submerged
46. Add 13μL of 10X loading dye to the pooled 120μL of PCR products from step 44, pipette to mix
47. Load entire volume of pooled DNA split across 4 wells (∼33.25μL each), otherwise the volume will overflow the wells
48. Load 10μl of 100bp ladder and run gel at 70V for ∼1.5h or until loading dye has migrated one half to two thirds of the way down the gel
49. Turn off the gel apparatus and remove the gel to the UV transilluminator, you may allow it to cool for several minutes so the gel does not fog up the glass **(CRITICAL STEP) ensure the transilluminator surface has been decontaminated with bleach or DNAZap, cover the surface with plastic wrap to ensure there is no contamination between gels**
49. Using protective sleeves and UV filtered goggles excise the bands of interest (should be around 300bp mark) from using the scalpel blade and place in pre-labeled and pre-weighed screw cap tubes
50. Repeat this for all 4 wells in the gel, these will be combined later in the gel extraction but the initial volume is too large to fit in a single tube
51. Weigh the tubes again to calculate the amount of agarose that must be dissolved, add 3 volumes of QIAGEN buffer QG from QIAquick Gel extraction kit (i.e. for 100mg of agarose, add 300μL buffer QG)
52. Incubate at 50 °C for 10 min or until the gel dissolves, mix by vortexing every 2-3 min to dissolve the gel
53. Color of mixture should be yellow after gel has dissolved completely. If not, add 10μL 3 M sodium-acetate pH 5.0 and mix.
54. Add 1 gel volume of 2-propanol and mix. (example: for 100 mg fragment, add 100μL isopropanol).
55. Apply 750μL of the mixture to QIAquick spin column from QIAquick Gel extraction kit. Spin 1 min at 13,000 rpm. Discard the flow-through, load another 750μL, and spin again at 13,000 rpm. Repeat this step until the whole QG / 2-propropanol / sample mix is used, from both tubes / fragments. Discard flow-through.
56. Add 500μL Buffer QG. Spin 1 min at 13,000rpm. Discard flow-through.
57. Add 750μL Buffer PE from QIAquick Gel extraction kit, keep for 2-5 min, and spin for 1 min at 13,000 rpm.
58. Discard flow-through, wipe down the column with a lint-free Kimwipe and put it in a new collection tube.
59. Incubate with the cap open for 5 min at room temperature.
60. Spin 2 min at 13,000 rpm and discard the flow-through. At this stage change the gloves.
61. Put column in 1.7 mL microcentrifuge tube and add 50μL Buffer EB from QIAquick Gel extraction kit or water (pH 7-8.5).
62. Incubate for 5 min at room temp.
63. Spin 1 min at 13,000 rpm. Briefly label the 1.7 mL snap-cap tube and discard column. Depending on your experiment, adjust volume of EB Buffer.
64. Transfer the eluted DNA to a pre-labeled 1.5 mL screw-cap microcentrifuge tube.
65. Quantify product by repeating step 30 to ensure adequate concentration for library sequencing
66. Perform paired-end sequencing with kit type: v3 25M and read length: 2×300bp on libraries from step 64 on the Illumina MiSeq platform

##### Bioinformatics Analysis

**(CRITICAL STEP) this analysis pipline utilizes Python/3.7.0, Biopython/1.75, Mafft/2.6, Trimmomatic/0.36, FLASH/1.2.11. All analyses are performed using the computing cluster provided by Digital Research Alliance of Canada (formerly called Compute Canada), with Slurm and StdEnv/2018.3 module. Ensure all dependencies are installed. Alternatively, you may perform trimming and read combining manually and skip to step 70. You will need access to a linux based environment.**

67. Ensure all data is in raw FASTQ format named with the suffix “_R1_001.fastq” and “_R2_001.fastq” for the forward and reverse reads respectively
68. Pull repository from Github (https://github.com/BoxeRZL/UMI-DloopSeq) and update the environment variables by editing the configuration file ‘config.RX’ :

~~~
runName=RX*
dataLocation=RX-sequencingData**
laneListFile=RX-laneList.txt***
trimThreshold=30****
FlashThreshold=5*****
outputFileName=RX-mutations
~~~

* ‘RX’ is the name of the run (i.e. R11; edit it to the name you need)

**Data location is the folder where row FASTQ files are (i.e. R11-sequencingData)

***Lane list file is a test files containing a list of all lane names (i.e. R11-laneList.txt). It is for automated analysis to proceed all FASTQ files (i.e. “_R1_001.fastq” and “_R2_001.fastq”)

****Trim threshold is the minimum quality score that is acceptable during trimming.30 is the default. The threshold may be increased for more stringency or decreased if the sequencing data is of low quality (see the documentation of Trimmomatic for details, http://www.usadellab.org/cms/uploads/supplementary/TrimmomaFc/TrimmomaFcManual_V0.32.pdf )

*****Flash threshold is the minimum length of overlap required between the forward and reverse reads. 10 is the default. Various flash thresholds were tested by using our data and 5 is the optimized threshold. It may be edited if necessary. (see the documentation of FLASH for details, http://ccb.jhu.edu/software/FLASH/MANUAL )

**(CRITICAL STEP) We recommend using a computing cluster for large projects**

##### Reads trimming and combining

69. Trim sequencing adapters of Illumina and filter out reads with poor-quality bases and then combine paired reads on a minimum length of overlap. The default settings are specified in the configuration file ‘config.fileRX’. The reads surviving of trimming and combining are counts in percentages for quality control and can be found in an output file labeled as ‘preprocessingQC.csv’ in the data location folder (see Box 2 for interpretations of quality control parameters and considerations). Template documents to review lane quality see supplement file 1

~~~
bash 01preprocessing-QC-automated.sh ./config.fileRX
**Dependencies of processing** 01preprocessing-QC-automated.sh:
01PID-dloop-preprocessing-QC.sh
01preprocessing-QC-automated.sh
~~~

##### UMI barcoded reads grouping and initial alignment

70. Group barcoded reads with same UMI into a family. All UMI barcodes with the presence of less than two reads are not grouped and are appended to an output file labeled as ‘ungrouped.txt’. All reads without a recognizable UMI are listed in an output file labeled as ‘mutated.txt’. An initial alignment is processed between reads within a family to generate a family consensus which is appended to an output file labeled as ‘Consensus.aln’. The number of reads in each family is calculated into an output file labeled as ’groupsize.csv’ for subsequent analysis steps.

~~~
bash 02automated-wrapper-R9patch-clean.sh ./config.fileRX
**Dependencies of processing** 02automated-wrapper-R9patch-clean.sh:
02sbatch-MIDgrouping-R9patch-clean.sh # parameters of sbatch are specified here and can be edited as necessary
02MIDgrouping-inUse-R9patch-clean.py
02parseGroups-WTadjusted-R9patch-clean.py
~~~

##### Summarization of the number of reads among UMI families

71. Summarize the number of constituent reads among all families in an output file labeled as ‘groupSizesSummary.csv’. This is used to indicate whether there was sufficient amplification for quality control.

~~~
bash 03automated-groupSizeParse.sh ./config.fileRX
**Dependencies of processing** 03automated-groupSizeParse.sh:
03parseGroupSizes.py
~~~

**(CRITICAL STEP) if there is insufficient amplification see Box 2 for troubleshooting and adjust the volumes in step 33 and 34 accordingly.**

**(CRITICAL STEP) If there is insufficient amplification for certain specimens, rerun step 12 - step 32 for the specimens. If there continues to be insufficient amplification, do not dilute AMPure purified product and continue with protocol.**

##### Low frequency mutations calling

72. Consensuses of UMI families are aligned within each specimen and referred to the cambridge reference sequence (NCBI ID: NC_012920.1, https://www.ncbi.nlm.nih.gov/nuccore/251831106). For this protocol, positions of the reference sequence from 16559 to 279 were used. To parse low frequency mutations, mutations are accepted from the consensuses that have sequences with the presence above 75% reads and were generated from UMI families with above 5 reads.

~~~
bash 04automated-writeMutations-clean.sh ./config.fileRX
bash 04merge-worker-outputs.sh
**Dependencies of processing** 04automated-writeMutations-clean.sh:
04sbatch-writeMutations-clean.sh
04writeMutations-multiprocessing-clean.py
~~~

**(CRITICAL STEP) any sequences that have more than 15 gaps or mismatches in a pairwise alignment with the reference sequence is considered contamination and filtered out of the analysis. All contaminated reads are appended into an output labeled as ‘contamination.txt’. The number of contaminations is considered as a quality control consideration (Box 2).**

73. Estimate mutation burden based on the equation (Figure S2).

###### Box 1

**Protocol adjustments for low mtDNA copy number specimens**

Some specimens (i.e. sorted cell subsets from biobanked PBMCs) may have less than 500 copies/μL of mtDNA. In these cases, you may:

1. Decrease the amount of water in the mastermix in step 12 of the protocol

| Reagent | Volume (μL) |
| --- | --- |
| UltraPure water | 14.75 |
| 10X PfuUltra II Reaction Buffer | 2.5 |
| dNTP (25mM) | 0.25 |
| PfuUltra II polymerase | 0.5 |

And in step 13 increase the amount of DNA to 5μL

2. Perform the PCR reaction (step 33 - step 35) in duplicate for each specimen, then pool duplicates after reaction

**NOTE:** One concern when sequencing specimens with low levels of mtDNA is increased variability in the number of amplicons from each specimen after PCR. If several specimens contribute disproportionately less amplicons than others in the final library preparation submission, they may have low numbers of families after sequencing. To mitigate this, the following protocol alterations can be made:

3. In step 40: prepare a gel casting tray with two clear gel combs with 7 teeth, 4mm wide, and 6mm deep. Place one comb at the top of the gel and one comb in the middle of the gel.
4. Do not pool specimens (skip step 44).
5. In step 46: add 6μL of 10X loading dye to 60μL of PCR product from each specimen, pipette to mix.
6. In step 47: for each specimen, load entire volume of PCR product into each well **(CRITICAL STEP) ensure that each gel only contains specimen from one lane to avoid contamination between lanes**
7. In step 50 - step 65: extract DNA from each well separately.
8. After step 65, quantify purified product by repeating step 30. For each lane, pool specimens so as to normalize the amount of amplicons from each specimen in the final pool. Run each pool through QIAquick column and elute in 50μL Buffer EB or water (repeat steps 56 - steps 65).

###### Box 2

**Quality control parameters and considerations**

**Box 2: Quality control parameters and considerations**
| Parameter | Scope | Recommended threshold | Troubleshooting |
| --- | --- | --- | --- |
| **Reads passing trimming** | Per run | 90% with both forward and reverse reads retained | If quality scores are poor overall, may decrease the quality threshold, we recommend checking the run parameters as they may need adjustment |
| **Reads passing forward/reverse combination** | Per run | 70% of read pairs combined | If the read quality is good at the beginning of the read but drops off in either the forward or reverse read, may decrease threshold but it is recommended to check the read quality (FastQC) |
| **Number of families formed** | Per sample | At least 200 families | Accepting samples with fewer families offers less granular data, to increase the number of amplicon copies used in step34 |
| **Number of contaminated sequences** | Per sample | <1% of families | May indicate contaminated working environment or DNA samples, if contaminated with nuclear DNA sequences revise primer specificity. |

## Author Contributions

ASZ, IG and HCFC designed the original procedure. ASZ, ZL, RD, LCC, HC and IG performed experiments used in assay development. ZL, RD and ASZ developed the bioinformaFcs analysis pipeline. ZL, RD and LCC provided analysis and data visualizaFon. LCC and HC performed opFmizaFon experiments. ZL, RD and LC wrote the manuscript. All authors contributed edits and changes to the manuscript and approved the submitted version.

## Acknowledgments

We would like to thank the Sequencing and BioinformaFcs ConsorFum at the University of BriFsh Columbia, Vancouver BC, Canada. This work was partially supported through the following grants to HCFC: PJT-156007, HET-85515, and TCO-125269, from the Canadian Institutes of Health Research (CIHR). CTN-277 and CTN-291 from the CIHR Canadian HIV Trial Network (now CTN+), and a grant # 20004 from the Canadian Association for AIDS Research (CANFAR). ZESL was partially supported by a University of British Columbia Centre for Blood Research graduate scholarship.

## Supplement material

**Table S1:**
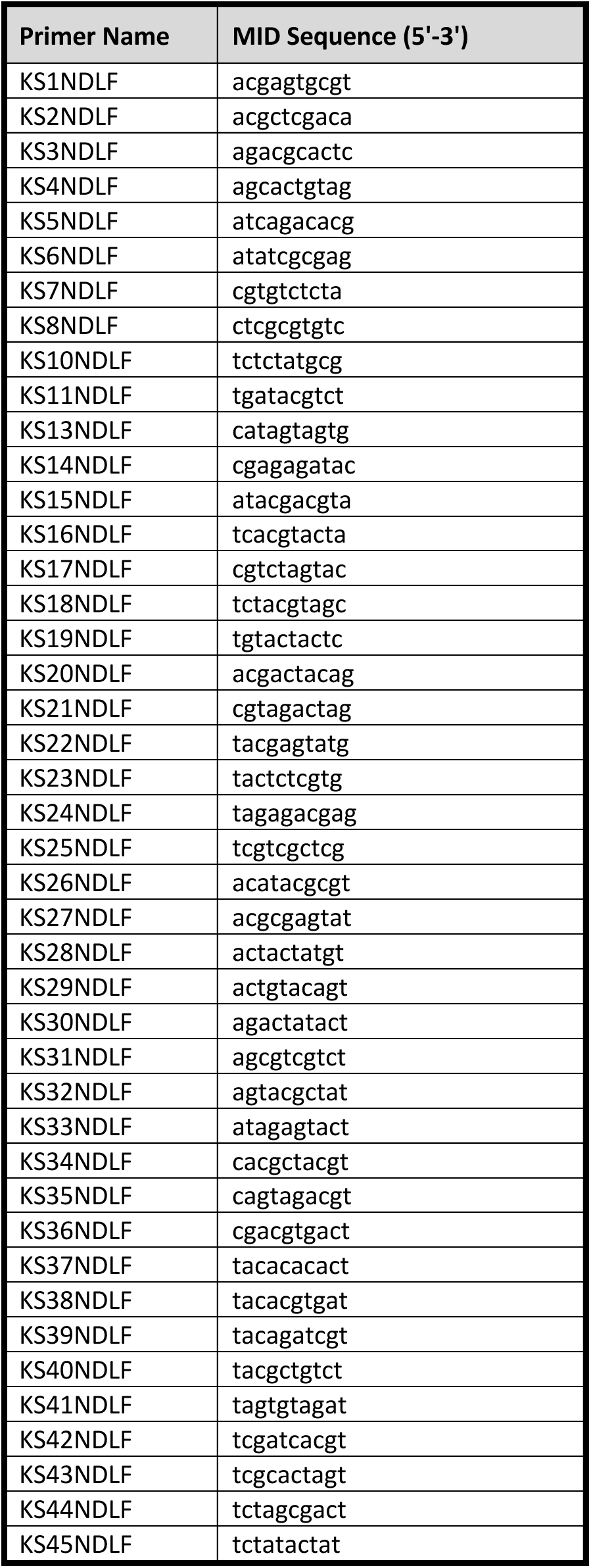
Extended list of MID sequences.

**Figure S1:**
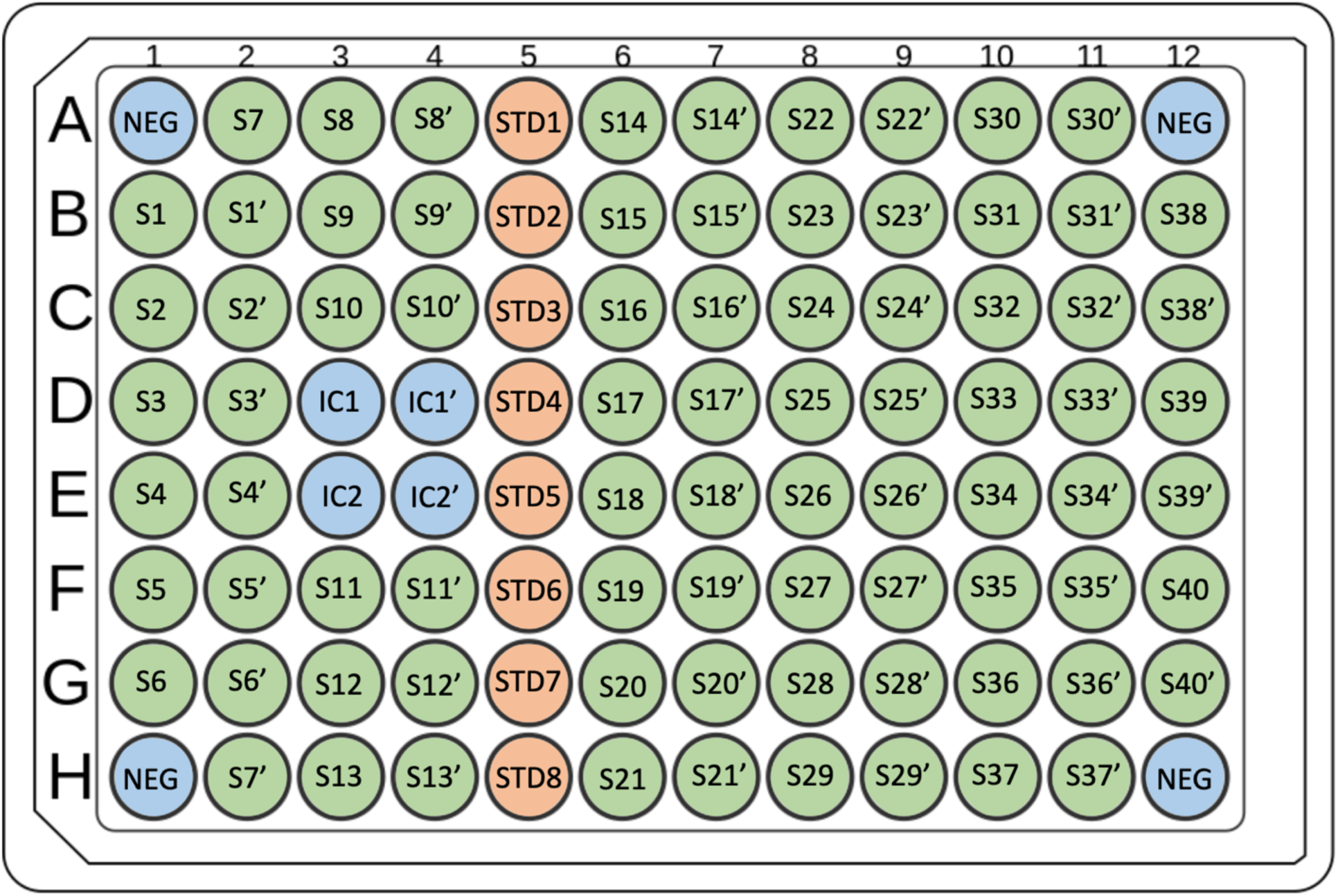
qPCR plate layout. Blue wells denote controls. NEG indicates negative controls, two should be PCR-grade H2O and two should be AE buffer for DNA elution and dilution. IC denotes internal controls of known value; one relatively high concentration and one relatively low concentration. These are assayed in duplicate, where ’ denotes the second well for that control. Green denotes specimens. These are assayed in duplicate, where ’ denotes the second well for that specimen. Orange denotes the standard curve. STD indicates standard dilutions arranged from highest concentration to lowest concentration.

**Figure S2:**
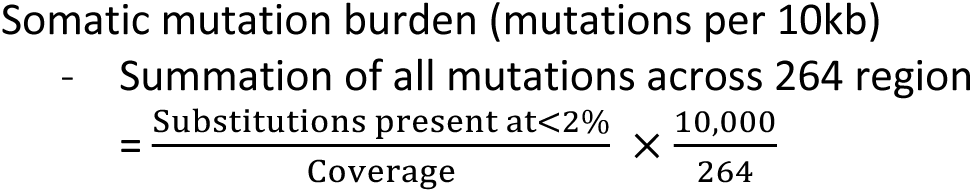
Equation of estimating overall mutation burden. The burden of substituted mutations is represented by mutations per 10kb. 264 bp is the length of the sequenced region utilized in this protocol.

